# Multicellular behavior enables cooperation in microbial cell aggregates

**DOI:** 10.1101/626481

**Authors:** Ali Ebrahimi, Julia Schwartzman, Otto X. Cordero

**Affiliations:** Department of Civil and Environmental Engineering, Massachusetts Institute of Technology, Cambridge, MA, USA 02139

## Abstract

During the degradation of biological materials such as biopolymers, extracellular enzymes liberate oligosaccharides that act as common goods and become available for all cells in the local neighborhood. This phenomenon can lead to cooperative growth, whereby cell-cell aggregation increases both the per-capita availability of resources and the per cell growth rate. However, aggregation can also have detrimental consequences for growth, as gradients form within aggregates limiting the resource accessibility. We used a computational model to show that high bacterial densities and high enzyme secretion rates restrict cooperation in aggregates larger than 10μm, due to the emergence of polymer and oligomer counter-gradients. We compared these predictions against experiments performed with two well-studied alginate degrading Vibrios, one of which displayed a strong density dependent growth. We observed that both strains can form large aggregates (<50μm), overcoming diffusion limitation by rearranging their internal structure. The non-cooperative, strong enzyme producer formed aggregates with internal channels that allowed exchange between the bulk environment and the aggregate core, whereas the cooperative, weak enzyme producer formed dense aggregates that developed a hollow structure as they grew. These internal structures allowed cells to avoid overcrowded areas near the core, enabling the development of large cell aggregates. Our study shows that bacterial behavior can help overcome competition imposed by resource gradients within cell aggregates.

## Introduction

Microbes live in communities, that is, collectives with thousands of cells influencing each-others function and behavior. In many cases, the activity of the collective can give rise to emergent forms of behavior and spatial organization that enhance individual fitness. For instance, *Myxococcus xanthus* and *Dictiostelium discoideum* form multicellular groups to forage for food and differentiate into stalked fruiting bodies to propagate spores through the air (1). Lichen communities and microbial mats segregate into distinct layers (2–4), while multispecies oral biofilm communities, methane and ammonia oxidizing consortia, and nitrogen-fixing granules exhibit complex structure that may create proximity between taxa optimal for the growth of individuals (5–9). In these examples, the structure of the collective provides information about its function.

Polymer degraders are the primary recyclers of dead organic matter in the biosphere (10), and often form dense communities. Organisms in these collectives confront the problem of digesting objects that are insoluble and, in many cases, much larger than their membrane transporters. To deal with this problem, polymer-degrading bacteria have evolved strategies that turn the extracellular environment into a digester: large substrates are hydrolyzed by secreted enzymes and hydrolysis products captured before they are lost to diffusion, fluid flow, or competitors. Examples of this phenomenon are found in the degradation of proteins, mediated by secreted proteases (11), or complex carbohydrates, such as alginate, mediated by secreted alginate lyases (12,13). In general, primary producers store most carbon and energy in complex polymeric materials. Therefore, degradation via secreted hydrolytic enzymes plays a central role in the recycling of organic matter and the closing of element cycles in ecosystems.

One of the most interesting consequences of extracellular enzyme activity is the potential for so-called cooperative growth dynamics to emerge. Cooperative growth implies that the per-capita growth rate is not constant, as in classical exponential growth, but positively dependent on cell density (11). A simple explanation for these dynamics is that when populations obtain resources via extracellular enzymes they can facilitate their own growth: cells secrete enzymes, which release oligomers, which are used to produce cells, thereby creating a positive feedback loop. This effect is most pronounced when cells are able to recover only a small fraction of the oligomer released by their enzymes, so that having another cell in the neighborhood increases the local concentration of oligomers and the uptake rate (14). When these conditions are met, cells that form aggregates should grow at faster rates than those that remain isolated. This is because within cell aggregates the concentration of local hydrolysis products is higher, thus increasing the growth rate of oligomer-limited cells. Moreover, dense aggregates such as biofilms can increase the retention of hydrolysis products, further increasing local concentrations (15). Therefore, cooperative growth implies that, as a collective, populations can recover more of the product thereby increasing per capita growth rates.

Although aggregation allows cooperation between cells, close cell-cell packing can also become detrimental once competition for oligomers overrides the benefits of cooperation. Moreover, in a structured environment, growth processes can also lead to the formation of resources gradients (16–18), which create areas of low resource supply that emerge due to diffusion limitation and rapid consumption of nutrients. This suggests that at certain aggregate densities, the “returns” from aggregation diminish, thus limiting the size that aggregates can reach. Interestingly, microbes have the ability to respond to the stress created within overgrown aggregates by regulating their behavior (19) and rearranging their positions within the aggregate. However, the potential for dynamic rearrangements within cell polymer-degrading aggregates, and the consequences of rearrangement for aggregate structure and function have not yet been explored.

We seek here to understand the conditions that lead to beneficial aggregation among polymer-degrading cells and the potential for cell reorganization to enhance aggregate growth. To address these questions, we developed an agent-based physical model of self-organized aggregates formed during growth of microbial populations on a soluble polymer. We used this model to explore how the physiochemistry of aggregates regulates growth and efficiency of up-taking break down products in this geometry. We tested some of the predictions of our model using alginate-degrading isolates of the marine microbe *Vibrio splendidus* (12,20–23). Alginate-degrading microbes secrete different amounts of hydrolytic enzyme and have different requirements for cooperation, prompting us to ask how bacterial behavior in the population alters the structure of aggregates and the efficiency with which alginate is consumed by cells. We find that the benefit of aggregation is negatively related to the hydrolytic power of individual cells, and that novel aggregate forms, such as “channeling” and “mixing” emerge by rearrangement of cells, as design solutions that maximize carbon use efficiency.

## Results

In our model, as in our experiments, a cell aggregate is immersed in a polymer solution. The polymer (soluble alginate) can penetrate the aggregate by diffusion, where it interacts with enzymes (alginate lyases) that hydrolyze it, releasing oligomers that can be transported into cells (Figure 1A). Oligomers can diffuse and be consumed by cells based on Monod kinetics (Figure 1B). The interplay between these processes gives rise to spatial gradients that can influence cell growth and depend on aggregate density and size. To study the impact of aggregate density on the growth rate of individuals within an aggregate, we fixed the aggregate radius to 20 μm and measured the mean rate by which individual cells take up oligomers, as a function of increasing bacterial density (Figure 1C). In our model, soluble polymers are assumed to be at a concentration of 0.1 mg/L. Alginate lyase activity is assumed to be diffusion limited, as can happen if the enzyme is tethered to the cell membrane (12,24) or trapped in extracellular matrix (15).

**Figure 1.**
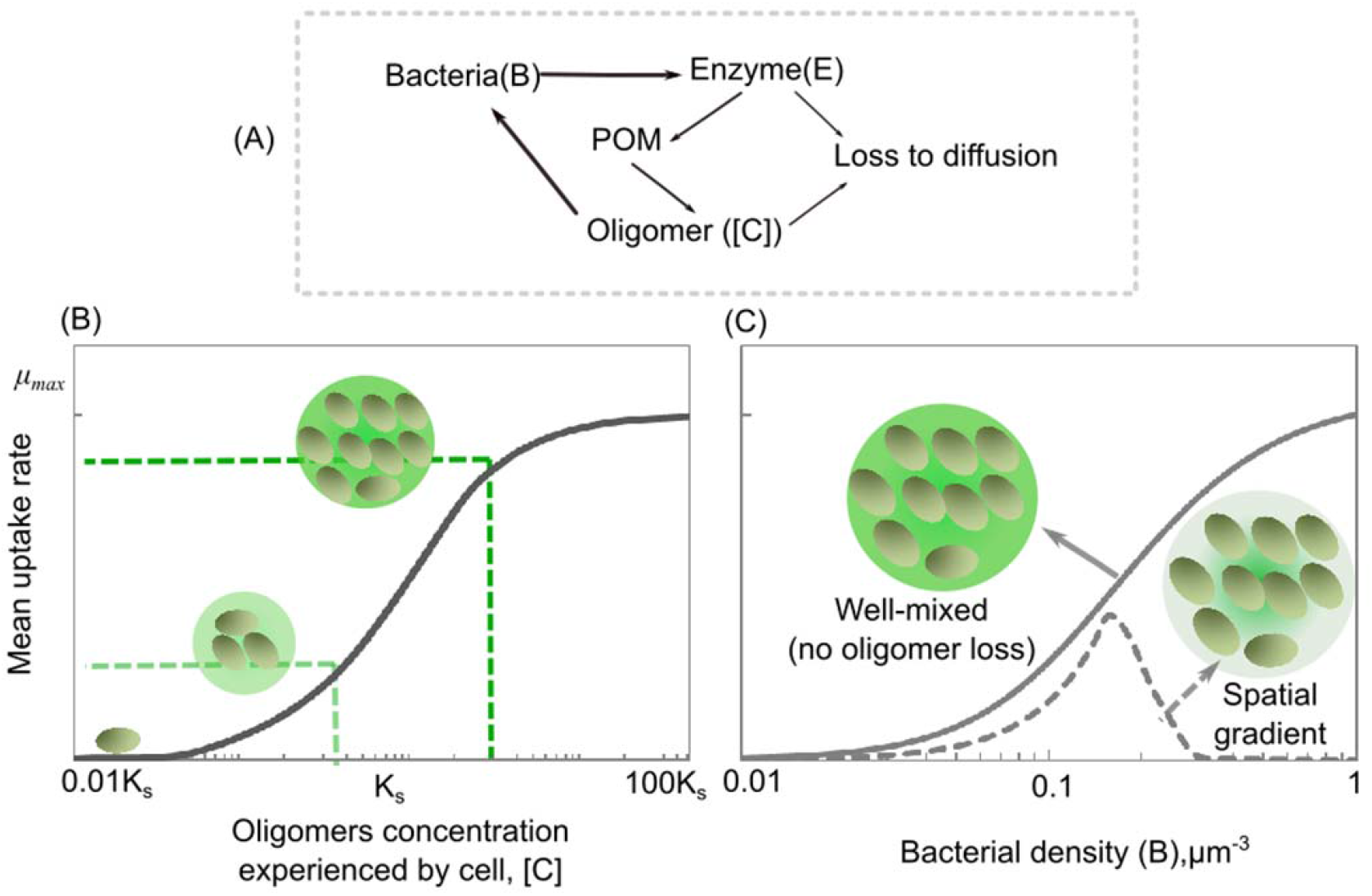
Bacterial aggregation in polymeric substrates leads to cooperative growth and enhances per capita uptake rate. (A) Conceptual representation of our model: Bacterial cells secrete enzymes that break down polymers, releasing oligomers which are taken up by bacterial cells, thus closing the loop. Enzymes and oligomers can be lost to diffusion (B) The mean cellular uptake rate is a function of oligomer concentration, which follows Monod kinetics. A schematic representation of the effects of initial cell numbers on oligomer concentration and its corresponding uptake rates are shown. (C) Mean uptake rate by bacterial cells are simulated as a function of initial cell density within a spherical aggregate of diameter 20 μm 20 h after inoculation. The simulation results are shown for a well-mixed closed system with no oligomer loss (solid line) and for an unmixed aggregate where diffusion is the dominant transport mechanism that allows for the loss of oligomers to the bulk environment at the aggregate periphery (dashed line). At low bacterial densities the aggregate behaves as a well-mixed reactor of size 20μm, indicating the absence of gradients.

### The balance between cooperation and competition

The results revealed an optimal cell density within the aggregate that maximized mean per capita growth (Figure 1C). At densities below the optimum the per-capita growth rate increased with cell density. The positive density dependence observed below the optimal cell density is consistent with the benefit derived from taking up oligomers released by neighbors, which would otherwise get lost to diffusion. This means that in this condition cells cooperate by sharing the oligomers released by their enzymes. For this positive dependence between uptake (growth) and cell density to emerge, an increase in the oligomer concentration should increase the per-capita uptake rate. Because in our simulations the relationship between oligomer uptake rate and oligomer concentration followed Monod kinetics (Figure 1B), the conditions for positive density dependence were those where oligomer concentration was approximately equal to the affinity of cells to oligomers, *K_s_*. In contrast, when the concentration of oligomers was significantly higher than *K_s_* the uptake rate saturated, limiting the benefit of increasing cell density on carbon uptake. From this point on, an increase in cell density was detrimental for the per-capita uptake (Figure 1C). In these conditions, competition for substrate overrides the benefits of cooperation.

To better understand the impact of cell density on cell growth, we studied the emergence of spatial gradients within aggregates of fixed size (20 μm) and low initial cell density (0.2 μm^−3^) (Figure 2A). Our simulation results revealed that counter gradients of polymer (diffusing from the outside, Figure S1) and oligomer (diffusing from the inside), lead to a narrow range of positions along the radial axis for which cooperative growth took place (Figures 2B). Outside this narrow band, towards the core of the aggregate, cells quickly experienced high competition due to the high cell densities attained and the slow diffusion of oligomers towards the core (Figure 2C). Outside of the cooperative range, towards the periphery, most oligomers released were lost by diffusion to the bulk environment, limiting the growth of cells. This lead to a situation where cooperative cells are “sandwiched” in between cells starved by the losses imposed by diffusion and by competition (Figure 2BC). Our model shows that, contrary to intuition, the more public good the cells make available, the faster the transition to competition occurs. This is because the higher the activity of the secreted enzyme per capita, the less cells cooperation is needed before growth rate saturates. Moreover, increasing the amount of enzyme expressed by a growing cell in an aggregate only increased the amount of hydrolyzed product available at the periphery (see Figure S2 for model results with higher enzyme production rate). Based on these results, we predict that aggregation supports cooperative growth only for a narrow range of cell densities (number) and cells with low enzyme production rates (Figure S3).

**Figure 2.**
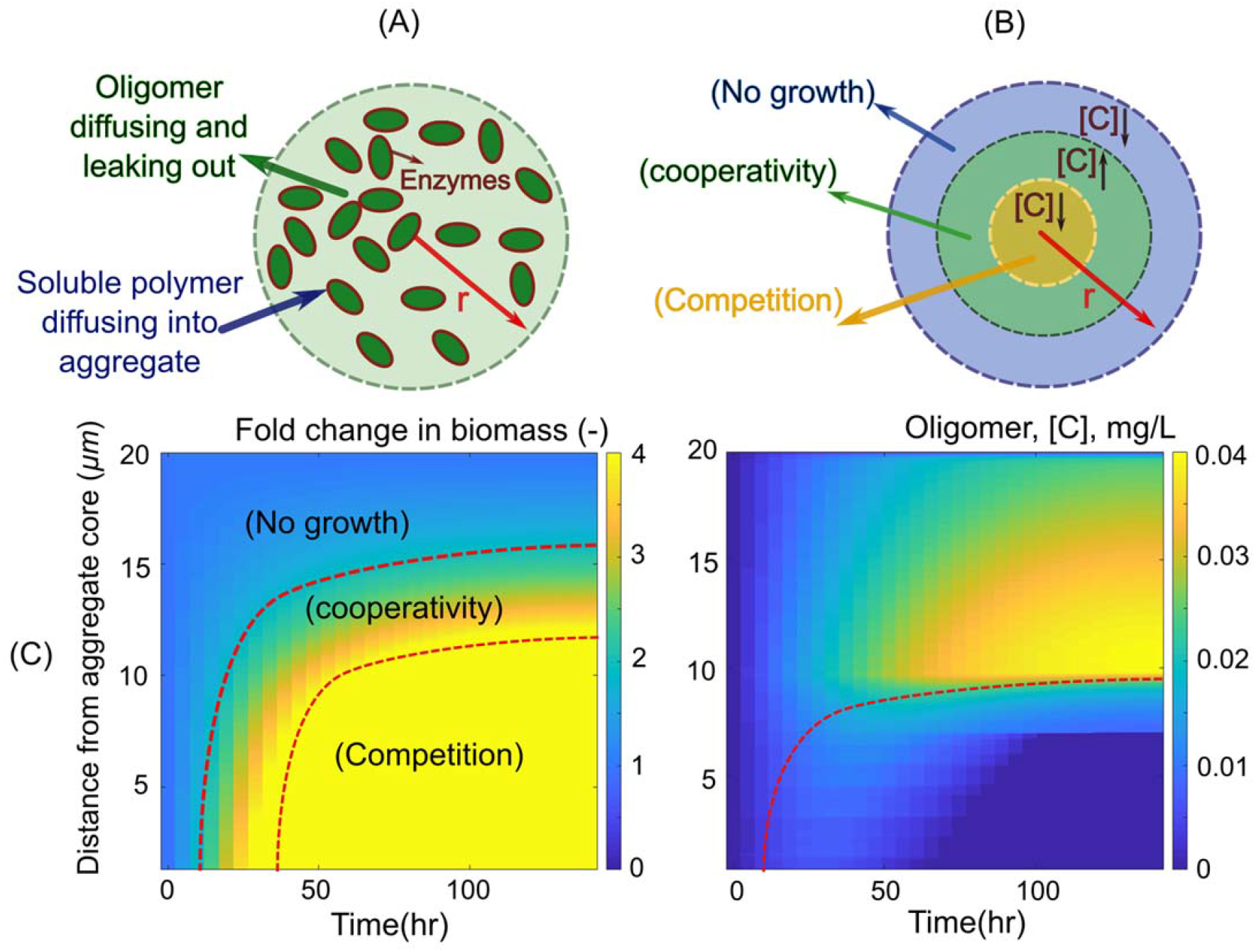
Spatial gradients restrict oligomer uptake by bacterial cells in aggregates. (A) Schematic representation of the individual based model developed for bacterial growth in an aggregate. The boundary condition for diffusion of oligomers is set to be zero at the aggregate periphery and soluble polymer is considered to be present at the aggregate surface at a constant concentration. Enzymes are assumed to be membrane bound with no diffusion. (B) Schematic representation of the narrow range of optimal oligomer concentration, predicted by the model, that allows for cooperative bacterial growth in aggregates. (C) The fold of change in biomass (left) and oligomer concentration (right) are shown along the aggregate radius over time, revealing the migration of a zone that optimizes biomass production and the emergence of competition within the core of the aggregate. The simulations are performed for an aggregate with constant radius of 20 μm and initial cell density of 0.2 μm^−3^. The enzyme production rate is assumed to be 0.02 hr^−1^.

### Multicellular behavior allows individuals to avoid competition

Our model is based on a number of simplifying assumptions, such as that cells are not able to move within the aggregate and remain uniformly packed. In reality, however, cells may have behavioral or physiological strategies to cope with the limits of diffusion within aggregates. To study possible aggregation strategies, we performed experiments in which we induced auto-aggregation of marine bacteria capable of degrading alginate. In particular, we focused on two strains of alginate-degrading *Vibrio splendidus*, 12B01 and 13B01. The two strains were chosen because they secrete different amounts of alginate lyase during growth on alginate polymer, leading to different hydrolysis kinetics (Figure 3A) (12). Moreover, we found that the weak enzyme broadcaster (12B01) exhibited positive density dependent growth when cultured with 0.1mg/L alginate as a sole carbon source. That is, we observed no growth when cells were inoculated below a threshold cell density, indicating that growth was positively dependent on population density (cooperative growth). By contrast, the strong enzyme broadcaster (13B01) did not display signs of density dependence, suggesting a much weaker tendency to cooperate. Both cultures contained cell aggregates, which appeared in absorbance readings as fluctuating measurements beginning in late exponential growth phase. To better understand these aggregates, we set up a culture system where we could periodically sub-sample cells as aggregate formation developed (Figure S4). To induce auto-aggregation, we incubated an initial population of 10^4^ CFU/mL in 70 mL 0.075% low-viscosity alginate in 250 mL shaking flasks (Figure S4). Consistent with our model predictions, we observed that the weak enzyme producer (12B01) formed aggregates with significantly higher bacterial density compared to the strong enzyme producer (13B01) (Figure 3B). The weak enzyme secretor, 12B01, formed densely packed aggregates, consistent with the notion that low hydrolysis rate per cell requires more cooperation. On the other hand, the loosely packed aggregates of 13B01 had a structure that resembled balls of crumpled paper, with folds and facets containing seemingly aligned cells (Figure 3B). Time lapses of 13B01 aggregates revealed that cells continuously rearranged themselves on the aggregates through frequent attachment and detachment (Movie S1). We speculated that this structure might represent a network of flow channels within the aggregates that promote the migration of detached cells, polymer, and hydrolysis product between clusters (Movie S2 and Figure S5). Simulations showed that channels larger than 3 μm in diameter improved uptake rates in large aggregates (>20 μm), while providing no benefit in small aggregates (<10 μm) (Figure 4). Therefore, the channeled structures formed by 13B01 may overcome diffusion limitation and facilitate exchange between the aggregate interior and the bulk environment.

**Figure 3.**
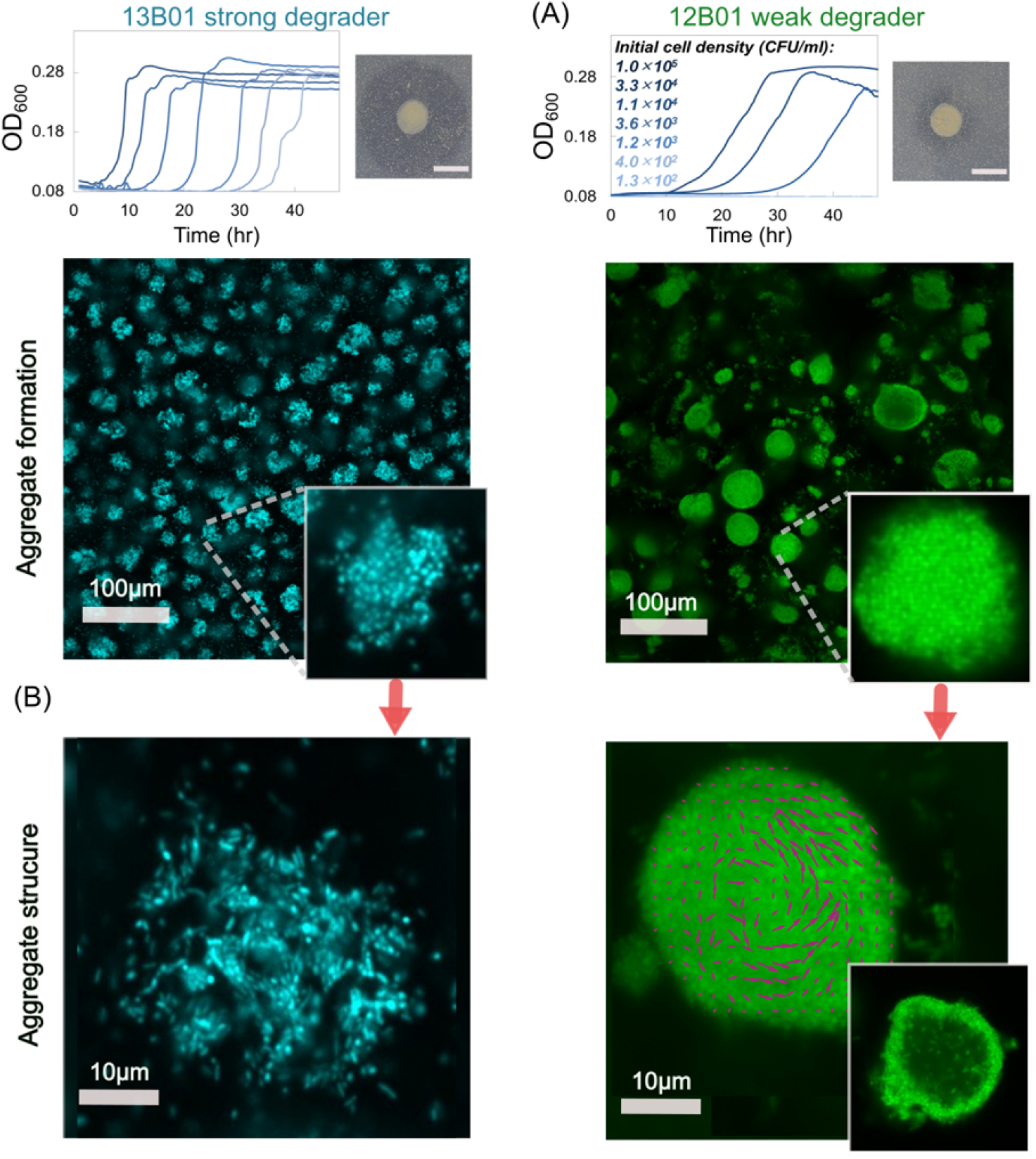
Bacterial enzymatic activity affects bacterial cooperative growth and resulting aggregate structure. (A) Density dependent growth for 12B01 and 13B01. Experiments were performed to measure the growth of these two strains in well-mixed liquid alginate as a function of the initial cell density. In the absence of density dependence, we expect growth curves to be delayed in proportion to the dilution factor, as in the case of 13B01. By contrast, if cells grow cooperatively, we expect delays that grow disproportionally with dilutions – or no growth whatsoever, as observed in 12B01. The broadcasted alginate lyase activity of 12B01 and 13B01 is shown on the right of each panel, measured as halos on cetylpyridinium chloride-treated alginate agar plates. The dark grey halo indicates the degree of broadcast alginate lyase activity originating from colonies of the two strains after 48 h of growth. (B) Autoaggregation of 13B01 and 12B01. The aggregate structure is shown as a maximum-intensity projection of 100 μm image stacks, and a cross-section taken in the middle of the aggregate is shown in the inset. Confocal images were taken after 24 h of incubation with 0.07%(w/w) alginate with shaking at 25 °C. A section taken at mid-plane (right) from 28 h old aggregates reveals that dense 12B01 aggregates undergo restructuring. Magenta vectors superimposed on 12B01 indicate the average velocity of cells over 5 frames (500 ms). The images were visualized at 40x magnification and 10x optical zoom was applied to highlight single aggregates.

**Figure 4.**
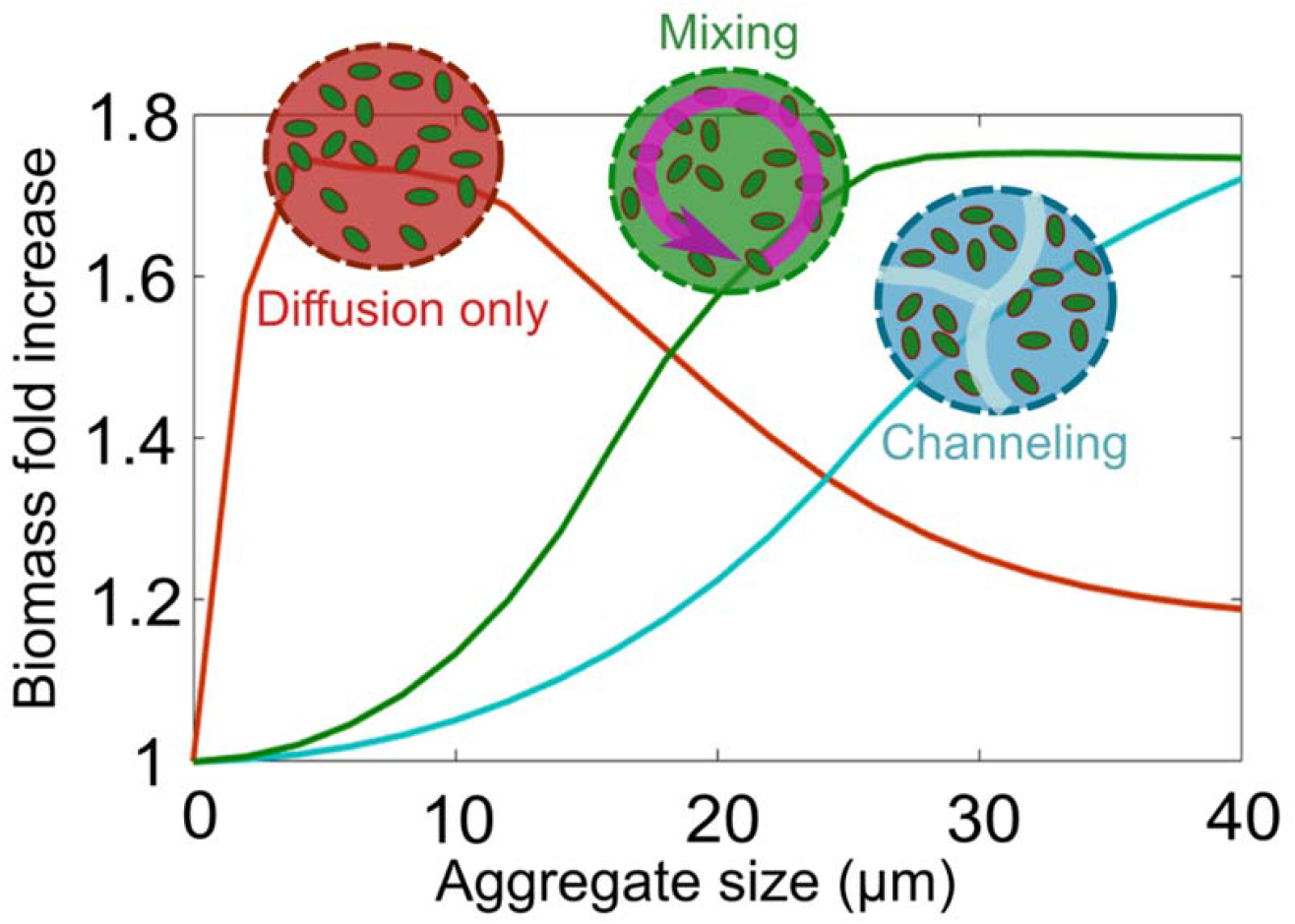
Bacterial reorganization within aggregates optimizes biomass accumulations in bacterial aggregates. Simulation results for biomass accumulation as a function of aggregate size are shown for the two dominant aggregation structures that we experimentally observed (Figure 3): mixing (green line), channeling (blue line). The reference scenario with no biological strategy, diffusion governed, is represented for comparison (red line). Aggregate size is shown at 24 h after inoculation. An initial cell density of 0.2 μm^−3^ and the enzyme production rate of 0.02 h^−1^ are assumed.

Despite the dense packing of the weak degrader 12B01, aggregates of this strain showed an average size of ~20 μm in diameter, with a maximum of more than 50 μm (Figure S6). This prompted us to hypothesize that the behavior of individual cells within the 12B01 aggregate might promote rearrangement along the radial axis, a strategy that would overcome the limits of diffusion and delay the onset of competition at the aggregate core. To measure behavior within the aggregate, we recorded timelapse images over periods of 10 min at a fixed cross section of the aggregate core. Surprisingly, we found that there was rapid mixing within the core of the dense aggregates formed by the weak degrader 12B01. This mixing emerged when the aggregates exceeded 15-20 μm in diameter (Figure 3B, Movie S3). The structures that emerged are reminiscent of the characteristic hollowing of surface attached biofilms that is observed during cell dispersal (25–27) (Movie S4). Complementing our observations with mathematical modeling of the flow patterns within the aggregate core demonstrated that mixing in the center of an aggregate can enhance resource acquisition in large aggregate sizes by homogenizing gradients (Figure 4). Collectively, these experimental observations and simulations suggest that hollowing and channel formation may be additional ways for cell aggregates to overcome size constraints imposed by resource gradients. Future work will be needed to better understand how different physiological attributes of individuals give rise to particular aggregate architectures, and how these forms shape the function of the collective in natural environments.

## Discussion

A known advantage of dense population clustering is that it allows cells to cooperate by sharing public goods. On complex substrates such as polymers, high cell densities locally concentrate enzymatic activity in a small area (“aggregates”), increasing the local concentration of hydrolyzed oligomer (28). In this study, we asked what physical and physiological constraints limit cooperative growth in aggregates, and how variation in the amount of enzymatic activity expressed by individual cells might give rise to structural variation of the collective. Our work demonstrates that the ability of cell aggregation to support cooperative growth is limited by counter gradients of polymer and oligomer, which are formed by diffusion and consumption (Figure 1). These counter gradients define a narrow zone within an aggregate where cooperative growth may be supported (Figure 2). Thus, hydrolysis and consumption of oligomers paired with substrate diffusion set limits on the size and the density of self-organized aggregates.

A key finding of our simulations and experiments is that the limitations of growth in colonies can be overcome by multi-cellular behaviors such as the formation of channeled aggregates or by mixing within the aggregate core (Figure 3B and Movie S2-S3). Such behavioral traits prevent the formation of gradients and the rise of competitive dynamics within aggregates. In particular, we observed that physiological variation in enzyme expression is correlated to two distinct strategies that our simulations predict mitigate competitive growth dynamics. In particular, growth dynamics in crumpled paper aggregates, formed by cells that express high levels of alginate lyase, are optimized by channels and by dispersal-driven exchange to and from the aggregate. In contrast, in densely packed aggregates, formed by cells that express low levels of alginate lyase, cooperative growth can be sustained if cellular motility mixes the center of densely packed aggregates (Movie S1) and forms a hollow interior (Figure 3B). While similar “hollowing” has previously been reported in the context of the seeding dispersal of mature surface-associated biofilms (29,30), our work suggest that mixing is a strategy to avoid competition at the aggregate core in the context of polymer degrading colonies.

Despite the importance of cell-cell aggregation in natural environments (31,32), and a molecular understanding of the mechanisms by which diverse microbial taxa adhere to surfaces (33) and each other (7), fundamental gaps remain to understand how emergent structures arise from the interactions between individual cells in a collective, and how such forms reflect the collective function of populations. Future work should focus on revealing the underlying mechanisms that give rise to emergent structures, such as phenotypic differentiations (34,35) motility (36), quorum signaling (37), and other physiological processes that support aggregate formation.

## Methods

### Mathematical modeling of bacterial aggregates

The mathematical model describes individual cell activity within spherical aggregates in the presence of radial chemical gradients. We developed an agent-based model to quantify single cell interactions with polymeric substrates including enzyme secretion, uptake of breakdown products, growth and division. The polymeric substrates diffuse into the aggregate from the aggregate periphery. The model assumes well-mixed conditions with no-accumulation of chemicals in the bulk environment (mimicking an open system) and thus the concentration of polymer and oligomer at the aggregate periphery is modeled as a constant concentration. The oligomer concentration is assumed to be zero at the periphery, so bacteria depend on polymer degradation within the aggregate to grow. Individual cells are uniformly distributed in the spherical domain and consume substrate and grow in response to local oligomer concentration. In our simulations cells are not motile. A more detailed description of the model is provided in the supplemental information.

### Bacterial aggregate formation experiments

Experiments were performed in shaken flasks. Each 250 mL flask contained 70 mL of a defined minimal media supplemented with 0.075% (w/v) low viscosity alginate soluble from brown algae (Sigma-Aldrich, A1112) and bacterial cells at an A_600_ of 1.0 were diluted 10^−3^. Flasks were incubated at 25 °C, shaking at 200 rpm. Flasks were incubated at 25 °C, shaking at 200 rpm. A low alginate concentration was used to avoid changing the viscosity of the medium and to reduce passive aggregate formation due to depletion interactions (38). To visualize bacterial aggregates and their spatial structures, 200 μL subsamples stained with the DNA-intercalating dye SYTO9 (Thermo Fisher, S34854) at a 1:285 dilution in 96-well plates with optically clear plastic bottoms (VWR 10062-900). To avoid evaporation from the wells, sterile self-adhesive sealing films were used to seal the 96-well plates. Additional experimental methods are described in the Supplemental information.

## Acknowledgments

We thank Martin Polz for the gift of strains 12B01 and 13B01, and Gabriel E. Leventhal, and Shaul Pollak for their comments and careful reading of the manuscript. We also thank all members of the Cordero lab and Fatima Hussain for their support and critical feedback. This project was supported by Simons Early Career Award 410104 and the Simons Collaboration: Principles of Microbial Ecosystems (PriME), award number 542395. A.E. acknowledges funding from Swiss National Science Foundation Grant P2EZP2 175128.

## Supplemental Information

### Movie Captions

Movie S1. Dispersal and attachment/detachment of cells around aggregates of 13B01. The frame rate is twice the real time.

Movie S2. Channeling strategy in aggregates from 13B01. The frame rate is twice the real time.

Movie S3. Mixing strategy in aggregates from 12B01. The frame rate is twice the real time.

Movie S4. Hollowing and dispersal strategy in large aggregates of 12B01. The frame rate is twice the real time.

## 1. Experimental Methods

### 1.1 Culture conditions

Bacterial strains used in this study were previously isolated by plating samples from size fractionated water samples [9–11]. For culturing from glycerol stocks, strains were streaked onto Marine Broth 2216 (Difco 279110) 1.5% agar (BD 214010) plates. Single colonies were picked and transferred to 3 mL liquid Marine Broth 2216 in an 18 mm test tube and incubated at 25°C, shaking at 200 rpm to establish seed cultures. Seed cultures were harvested after ~5 hours by centrifugation for 1 min at 10000 rpm (Eppendorf 5415D, Rotor F45-24-11). The supernatant was discarded and serial dilution of these cells were used to establish pre-cultures in pH 8.2 minimal medium supplemented with 20 mM glucose (see below for full recipe). Following overnight growth, cell density was measured in 1 cm cuvettes by absorbance measurements at 600 nm (A_600_) using a NanoDrop Spectrophotometer (ThermoFisher Scientific). Cultures with absorbance between 0.5-0.7 were used for experiments measuring positive-density dependent growth and aggregate formation.

### 1.2 Halo assays to measure alginate lyase activity

Alginate lyase activity was detected as described previously (1), with minor modification. Briefly, assay plates were prepared by adding filter-sterilized 62.5 mL 2.5 % low viscosity alginate to an autoclaved bottle of 500 mL marine broth containing 1.5 % agar (Difco) cooled to 50 °C. Plates were dried overnight, and 5 μL of cultured cells, prepared as described above, were spotted onto the plates. After 48 h of incubation at 50 °C, plates were flooded with a 10 % w/v solution of cetylpyridinium chloride (CP) and deionized water. CP s a cationic quatrinary ammonium compound that adsorbs to polymeric alginate, revealing undigested alginate remaining in the agar as an opaque haze. The plates were incubated with the CP for 20 minutes static at room temperature, at which point the solution was decanted. The plates were gently flooded with deionized water, so that colonies were not disturbed, and the water was exchanged 3 times to remove excess CP. Images of the colonies and ‘halos’ where polymeric alginate had been digested were obtained using a Cannon EOS Rebel T7.1 digital camera with a EF-S 35 mm f/2.8 IS STM Macro lens.

### 1.3 Minimal Medium

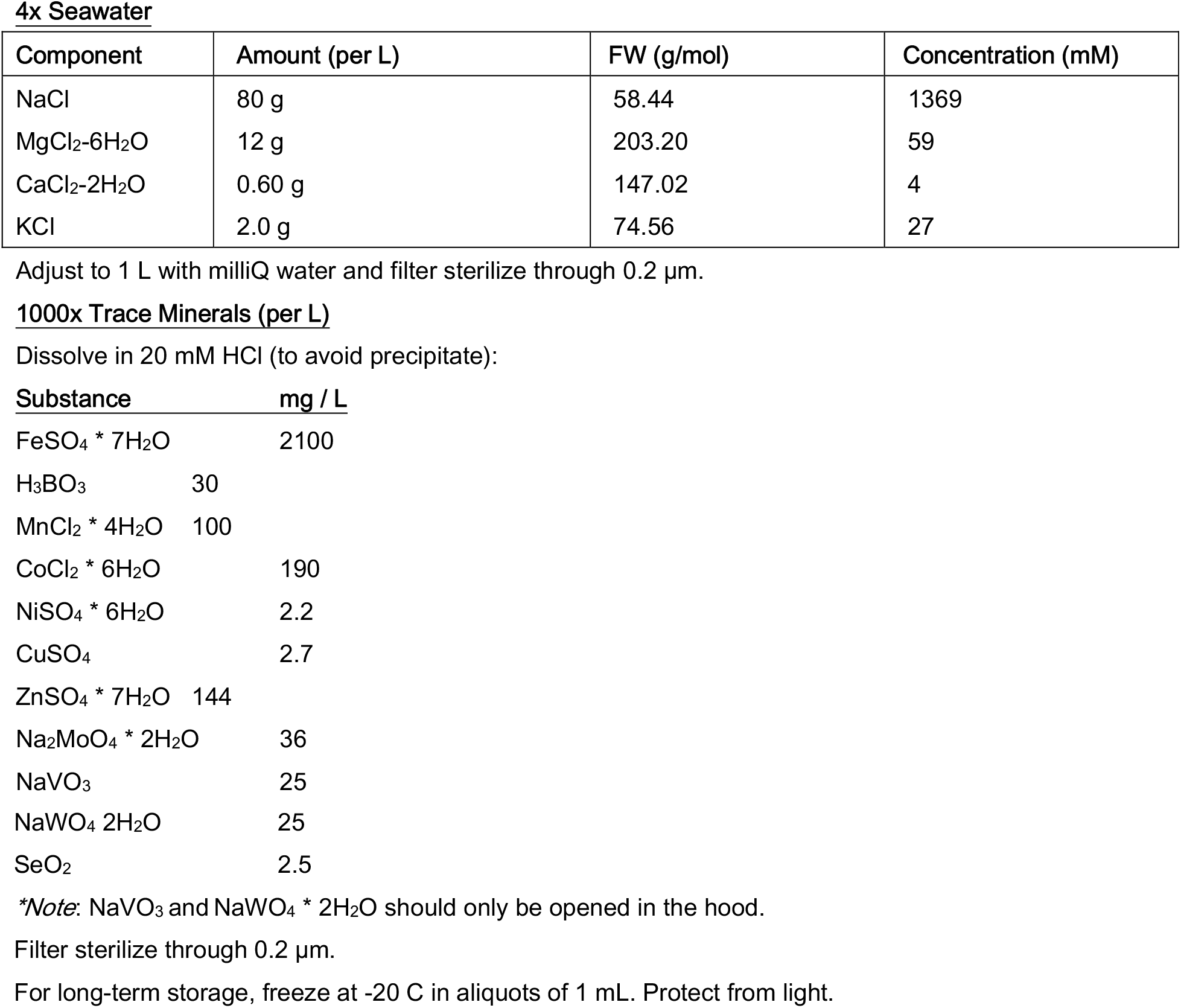

#### 1000x Vitamins

Dissolve in 10 mM MOPS, pH 7.2:

| **Substance** | **mg/L** |
| --- | --- |
| Riboflavin | 100 |
| D-Biotin | 30 |
| Thiamine hydrochloride | 100 |
| L-ascorbic acid | 100 |
| Ca-d- pantothenate | 100 |
| Folate | 100 |
| Nicotinate | 100 |
| 4-aminobenzoic acid | 100 |
| pyridoxine HCl | 100 |
| Lipoic acid | 100 |
| NAD | 100 |
| Thiamin pyrophosphate | 100 |
| Cyanocobalamin | 10 |

Titrate with a couple of drops of 5 M NaOH to avoid precipitate.

Filter sterilize through 0.2 μm.

Store at 4 C in the dark. Stable for a couple months.

For long-term storage, freeze at −20 C in 1 mL aliquots. Protect from light.

#### Nitrogen source

1 M ammonium chloride (100x)

Dissolve 2.14 g NH4Cl in 40 mL distilled water.

Filter sterilize through 0.2 μm.

#### Phosphorus source

0.5 M phosphate dibasic (500x)

Dissolve 2.84 g Na2HPO4 in 40 mL distilled water.

Filter sterilize through 0.2 μm.

#### Sulfur source

1 M sodium sulfate (1000x)

Dissolve 5.68 g Na_2_SO_4_ in 40 mL distilled water.

Filter sterilize through 0.2 μm.

#### HEPES buffer

1 M HEPES buffer (20x), pH 8.2

#### Per 100 mL

26.29 g HEPES sodium salt

Dissolve in 75 mL distilled water. Adjust pH to 8.2 with concentrated NaOH and constant stirring.

Bring final volume to 1 L with water. Filter sterilize through 0.2 μm. Store at 4 C.

#### Basic recipe for carbon-free medium (500 mL)

0.5 mL vitamins

0.5 mL trace metals

0.5 mL 1000x sodium sulfate

1.0 mL of 500x phosphate dibasic

5 mL of 100x ammonium chloride

25 mL of 20x HEPES buffer, pH 8.2

50 mL of 4x seawater

416.5 mL of sterile milliQ H_2_O

#### Glucose stock

1 M glucose (50x):

Dissolve 1.8 g D-glucose in 10 mL carbon-free minimal medium.

Filter sterilize through 0.2 μm. Store at 4 C.

#### Preparation of glucose minimal medium

Add 2 mL glucose stock to 98 mL carbon-free minimal medium.

#### Preparation of alginate minimal medium

0.1% Low viscosity alginate (Sigma A1112)

Add 0.1 g alginate to 100 mL sterile carbon-free minimal medium. Add slowly with stirring: heat <100 °C can help the polymer hydrate and dissolve.

Filter sterilize through 5 μm Sterivex filters.

### 1.4 Measurement of positive-density-dependent growth

To compare the ability of the two strains used in this study to grow on polymeric alginate, cells were grown as described above, then pelleted, and re-suspended at an optical density of 1.0. A multichannel pipette was used to transfer 150 μL of 1 g/L alginate minimal medium to the wells of a 96-well plate. These washed, concentrated cells were diluted 10^−3^ in 1 g/L alginate minimal medium, and 100 μL was added to the first column of a 96-well plate. A multichannel pipette was used to mix the wells in the column, and aliquot 50 μL of medium to the subsequent column. This established 3fold dilutions of the initial cell population. The 96-well plate was sealed with optically clear sealing tape, and the plate was incubated with double-orbital shaking at 25 C in a Tecan Spark plate reader. Absorbance (600 nm) was measured in 15-minute intervals. These measurements were used to test for the existence of a minimum cell density that supported growth.

### 1.5 Confocal microscopy and image processing

Microscopy was performed on micro-confocal high-content imaging system (ImageXpress Micro Confocal, Molecular Devices), using the 60 μm pinhole spinning disk mode. Fluorescent signal was visualized with a LED light source (Lumencore Spectra X light engine), bandpass filters (ex 482/35 nm em 538/40 nm dichroic 506 nm), at 40x magnification (Nikon Ph 2 S Plan Fluor ELWD ADM 0.60 NA cc 0-2 mm, correction collar set to 1.1), and a sCMOS detector (Andor Zyla). To visualize aggregates,100 μm image stacks sampled at 0.7 μm intervals were acquired in Z using MetaXpress software (version revision 31201). 3D-reconstruction of Z-stack images and movies were done using the maximum intensity projection feature of ImageJ distribution Fiji (ImageJ 1.52). 16 fields of view were acquired for each timepoint. Time-lapses were obtained under the same imaging conditions, with a 2-second interval. Aggregate cross-sectional areas were measured in MATLAB. Briefly, intensity-based thresholding was used to define a binary mask that distinguished aggregates from their surroundings. The area, and the average signal intensity (proportional to the average cell density) was obtained for each segmented aggregate. The mixing velocity within aggregates of 12B01 was analyzed using the Matlab (2015b) plugin PIVlab (version 1.4) (2–4). The average velocity was determined over 5 frames (10 sec), using the Fast-Fourier-Transform algorithm with 3 passes (window sizes 32, 16, and 8 pixels).

## 2. Mathematical modeling of bacterial aggregates

The mathematical model describes individual cell activity within conceptual spherical aggregates in presence of radial chemical gradients. We develop an agent-based model to quantify single cell interactions with polymeric substrates including enzyme secretion, uptake of breakdown products, growth and division. The polymeric substrates diffuse into the aggregate from the aggregate periphery. The model assumes well-mixed conditions with no-accumulation of chemicals in the bulk environment (mimicking an open system) and thus the concentration of polymer and oligomer at aggregate periphery is modeled as a constant concentration. Oligomer concentration is assumed to be zero at the periphery, so polymer degradation within the aggregate provides the major oligomer for bacterial activity. Individual cells are uniform randomly distributed in the spherical domain but consume substrate and grow in response to local chemical conditions of oligomer concentration. Note that no motility is considered for the individual cells. Detailed description of the model is provided in the supplementary.

The following represents derivation of mathematical formulations and procedures to model microbial growth, enzymatic activity and diffusion processes.

### 2.1 Growth and division

In this model, the growth kinetics of individual cells, *ν_s_* are dictated by Monod-type kinetics given as

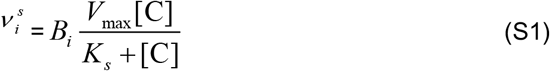

where *B* is the cell dry mass of an individual cell, *i* and *V*_max_ is the maximum substrate uptake and defined as: *V*_max_ = *μ*^max^ / *Y*_max_ (maximum specific growth rate / substrate conversion yield to biomass). *Ks* is the half saturation constant for oligomer s. *C_s_* is oligomer concentration and is assumed as the primary limiting substrate for the bacterial growth. We kept all other nutrients (e.g., oxygen, phosphate and nitrate) available at sufficient levels for microbial activity. Note that bacterial aggregates are often known to create anoxic microsites due to aerobic activity at their shells(5,6), however in our simulations the polymer concentration is kept low enough to avoid full oxygen depletion.

The actual biomass accumulation 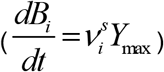 and maintenance of an individual cell 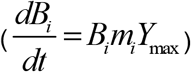 are linearly correlated with cell dry mass and therefore the new growth rate of individual cells (*μ_net_*) is given as:

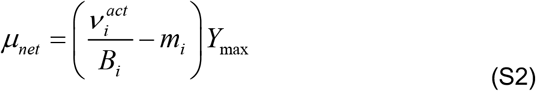

In the individual based model, each cell may double to two daughter cells when a threshold amount of substrate has been taken up (7,8). The minimum volume of the individual cell at the threshold for the division (*V^d,min^*) is given from the descriptive Donachie model (7,8)

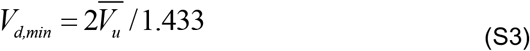

where 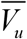 is median volume of the individual cell. The cell will divide into two identical (half-volume) cells if reaches *V_d,min_*. In the simulation, the cylindrical cells are assumed. Under limited nutrient conditions, the actual growth rate of an individual cell is restricted by actual available amount of substrate within the domain (see (9,10)). The biological parameters of the growth kinetics are provided in Table S1.

### 2.2 Enzymatic activity

Extracellular enzymes at individual cell level is modeled by assuming that the enzyme is bound to external membranes of the cells and no broadcasting or diffusion of enzymes is considered. Bacterial cells depolymerize the local alginate polymers and releases low molecular weight substrates. Enzyme level (*S_E,i_*) of a bacterial cell, *i* are assumed to be linearly correlated with cell biomass, *B_i_*:

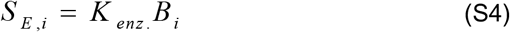

*K_enz_* is the fraction of biomass that is membrane bound enzyme and hydrolyzes polymeric substrates (enzyme production rate).

### 2.3 Oligomer and polymer diffusions along aggregate radius

The model explicitly simulates the diffusion oligomers and polymers within spherical aggregates. For oligomers, an absorbing condition (zero concentration) is assumed at external boundaries of the aggregates that simulates loss of oligomers to bulk environment. Diffusion of oligomers is modelled based on Fick’s law of diffusion. We modeled a spherical volume of a single aggregate and diffusion of oligomers is assumed to be only in radial direction. Reaction-diffusion equation is then numerically solved by finite-difference method:

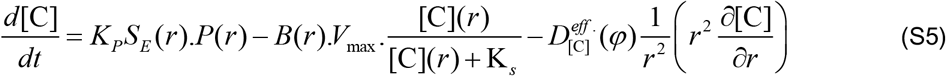

where [C] is the concentration of oligomers at radius, *r*. *K_p._* is polymer lability that defines how many grams of oligomers are released per gram of enzyme acting on the polymer surface per unit of time. *S_E_* is the enzyme amount and it is calculated from summing up the total enzymes of individual cells (*S_E,i_*, presence in the corresponding radius (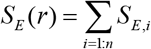, *n* is the number of individuals in the mesh grid at radius, *r*). *B* and *P* are the total biomass and polymer at radius, *r*, respectively. 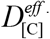 is the effective diffusion coefficient of oligomers. To take into account the effects of cell density on the diffusion coefficient, we treated the bacterial aggregate as a porous environment in which cell density modifies aggregate porosity, *ϕ*. Porosity of the aggregate is calculated based on number of cells, volume of individual cells and size of the aggregate. We applied Millington equation to provide the relationship between porosity and effective diffusion coefficients (11):

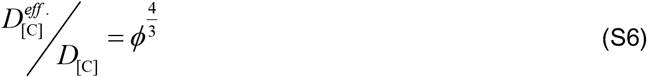

*D*_[C]_ is the diffusion coefficient of oligomers in bulk liquid.

Similar to oligomers, polymer diffusion is assumed to be only along the radial direction, as given:

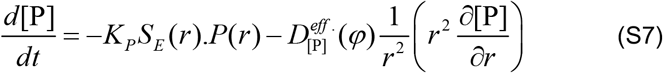

The physiological and chemical parameters used in the simulations are represented in Table S1.

#### Modeling biological strategies (mixing and aggregates with channels)

The main aggregate model is modified to implement biological responses on aggregate structure. In mixing strategy, the effective diffusion coefficients for both polymer and oligomers transport equations (Eq. S5 and S7) are enhanced to account for the effect of mixing on the transport.

To model aggregates with channels, we assumed that the channels are in equilibrium with the bulk environment and therefore the ultimate effect of the channeling is to subdivide the original aggregate structure into smaller individual sub-aggregates. We thus model the individual sub-aggregates as a separate unit and no interactions with other sub-aggregates are considered.

---

**Table S1.** Physiological and chemical parameters for microbial growth, metabolism and nutrient concentrations in the individual-based model.

| *Parameters* | *Values (Units)* |
| --- | --- |
| $\mu_{\max}$ : maximum growth rate ($hr^{-1}$) | 0.1<sup>*</sup> |
| $K_s$ : half saturation (mg/L) | 0.1<sup>*</sup> |
| $Y_{\max}$ : growth yield (gr dry mass/gr substrate) | 0.5<sup>*</sup> |
| cell maintenance | 0.1 $\mu_{\max}$<sup>f</sup> |
| cell size ($\mu m$) | 1<sup>f</sup> |
| $\rho$: cell density ($mg L^{-1}$) | $2.9 \times 10^5$<sup>f</sup> |
| $V_{1D}$ : cell velocity at bulk solution ($\mu m/s$) | 0<sup>*</sup> |
| $V_u$ : median cell volume (fl) | 0.4<sup>f</sup> |
| $K_p$ polymer lability ($hr^{-1}$) | 100<sup>ll</sup> |
| $D_{[C]}$ diffusion coefficient of oligomers in bulk liquid ($m^2 hr^{-1}$) | $2.4 \times 10^{-6}$ |
| $D_{[p]}$ Polymer diffusion coefficient in bulk liquid ($m^2 hr^{-1}$) | $1.7 \times 10^{-5}$ |

<sup>f</sup> (7)
<sup>τ</sup> (13)
<sup>ll</sup>Enzyme Database – BRENDA: mean observed value for alginate
<sup>\*</sup>Model assumption

## Supplemental Figures

**Figure S1.**
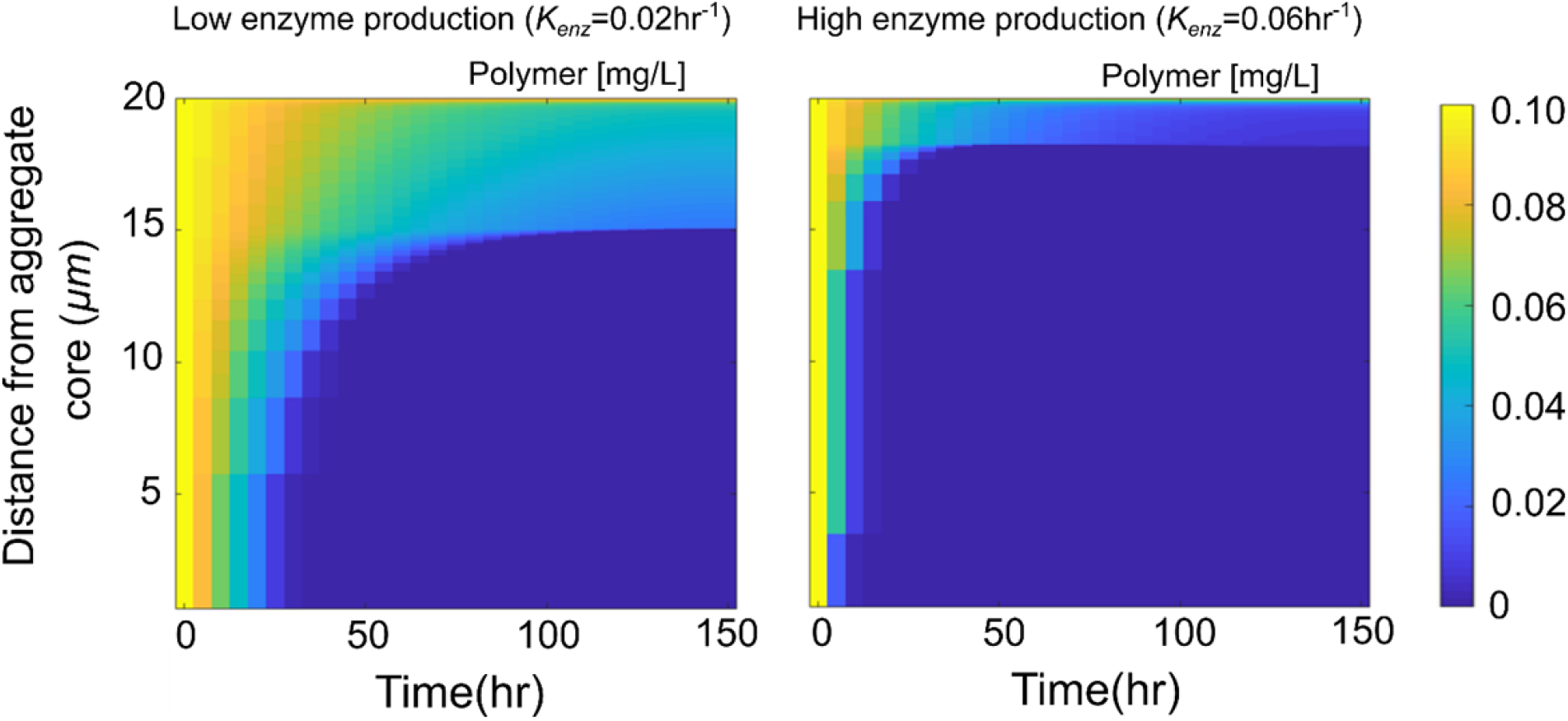
Spatial gradients of polymer concentration along aggregate radius over time for two enzyme production rates (*K_enz_*), respectively. The simulations are performed for an aggregate with constant radius of 20 μm and initial cell density of 0.2 μm^−3^.

**Figure S2.**
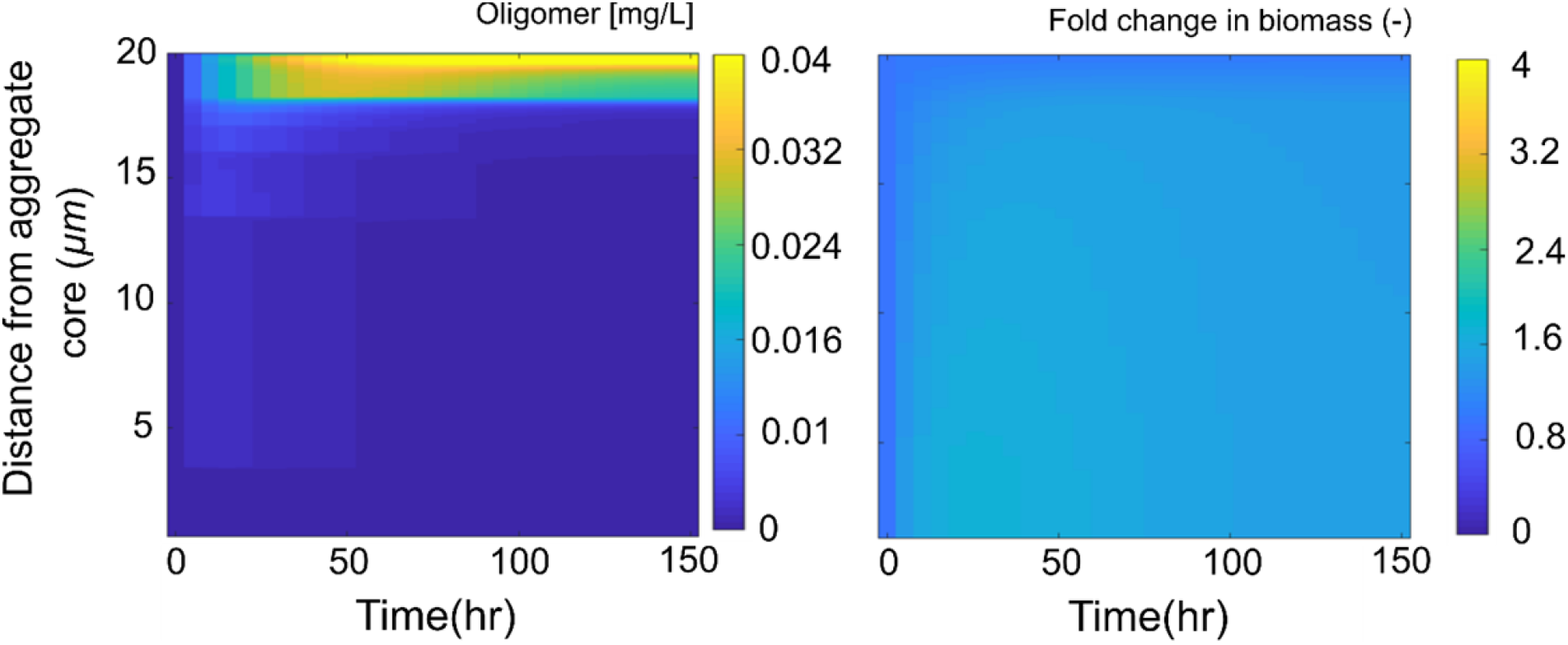
Oligomer concentration and fold change in biomass along the aggregate radius over time are shown for high enzyme production rates (*K_enz._*=0.06 *hr*^−1^). The simulations are performed for an aggregate with constant radius of 20 μm and initial cell density of 0.2 μm^−3^.

**Figure S3.**
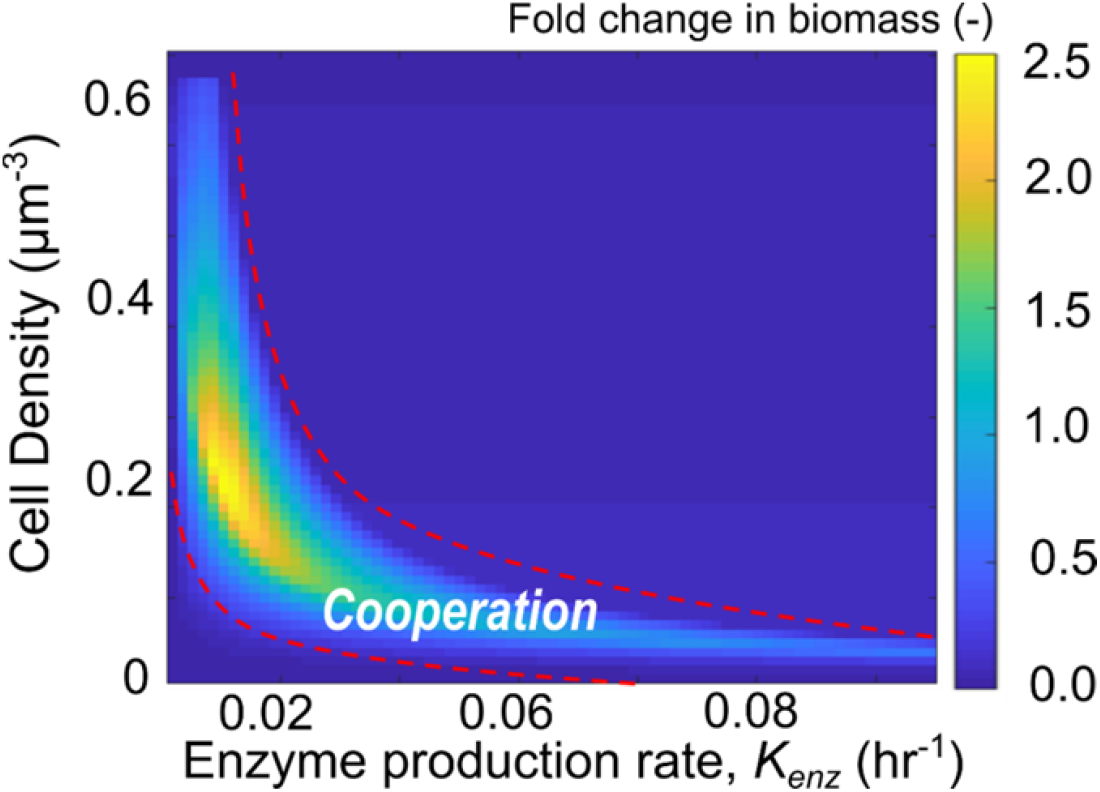
Fold change in biomass as a function of initial cell density and enzyme production rate are shown. Individual simulations are performed by assigning different values of cell density and enzyme production rates for an aggregate with a constant size (20 μm in radius). The results are shown for after 70 h. The fold change in biomass is averaged over the aggregate radius.

**Figure S4.**
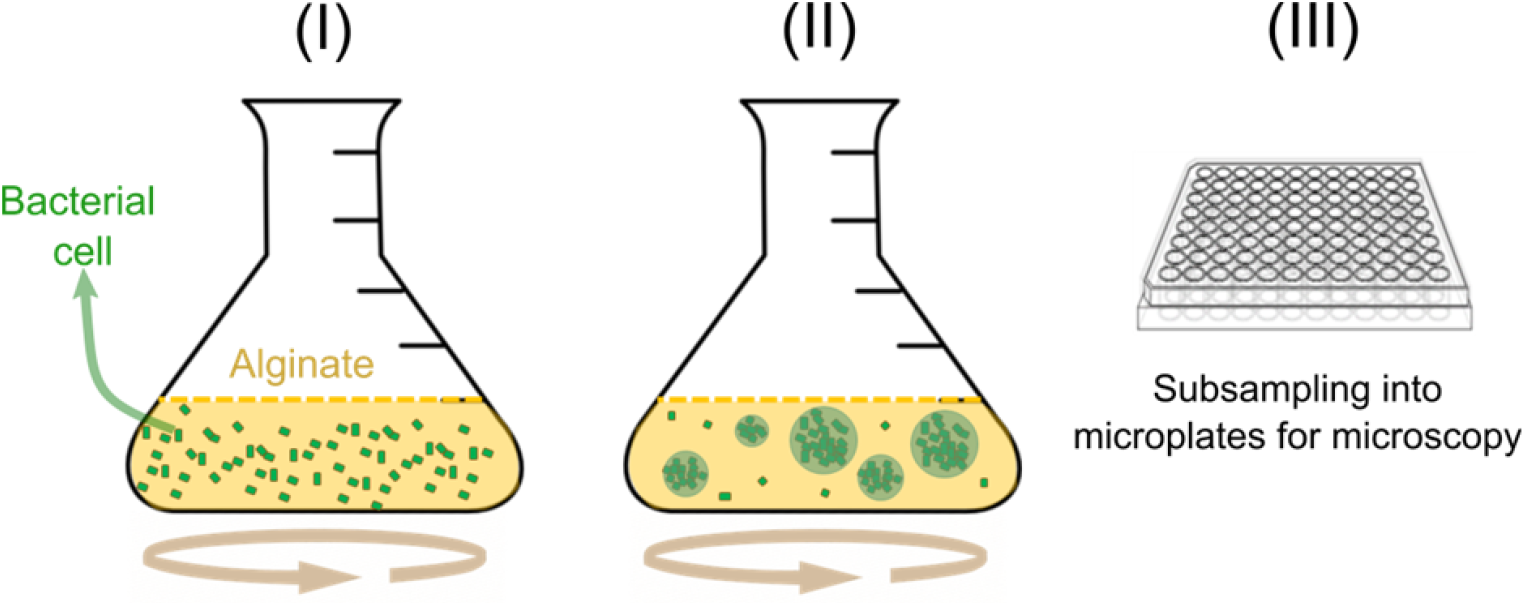
Experimental setup for monitoring aggregate formation and structure. Soluble low viscosity alginate is used as the main carbon source. Bacterial cells at low initial cell density (10^4^ CFU/mL) are used in shaking flasks with 70 mL solution. Subsamples of 200 μm from flasks are taken into 96 microplates for microscopy.

**Figure S5.**
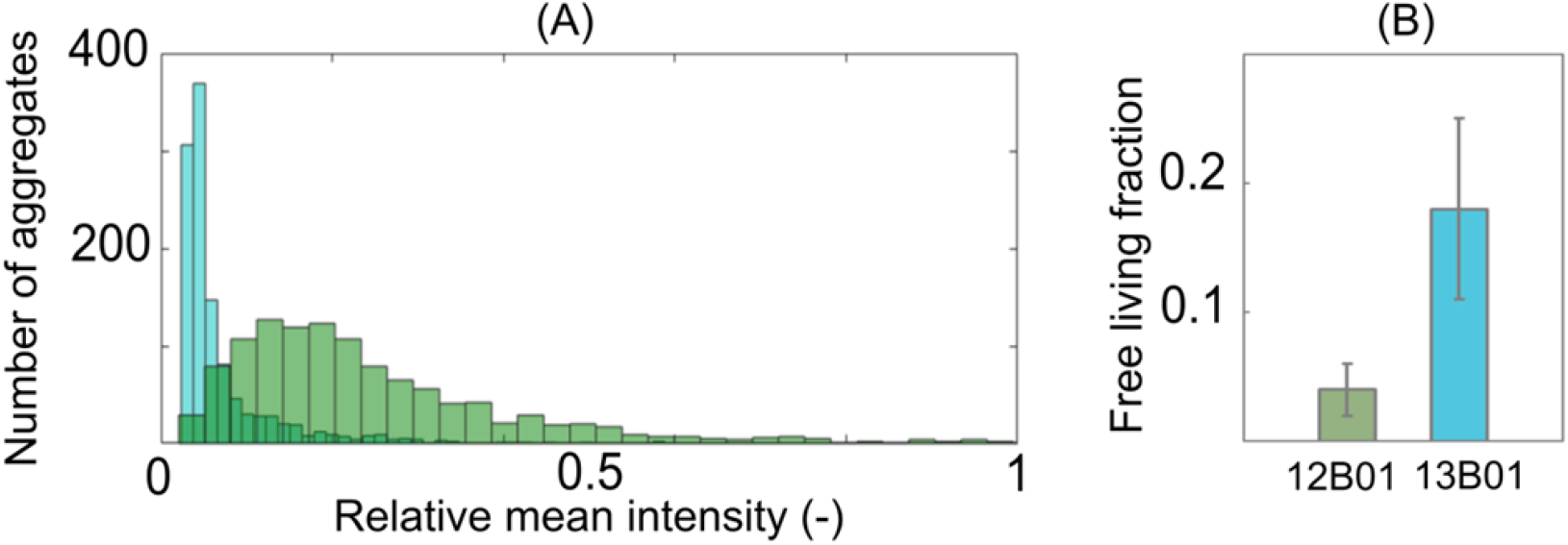
(A) Bacterial cell density distribution within aggregates, represented as relative florescent intensity. Over 1000 aggregates are analysed. (B) Number of free living cells is shown after 24 h from incubation.

**Figure S6.**
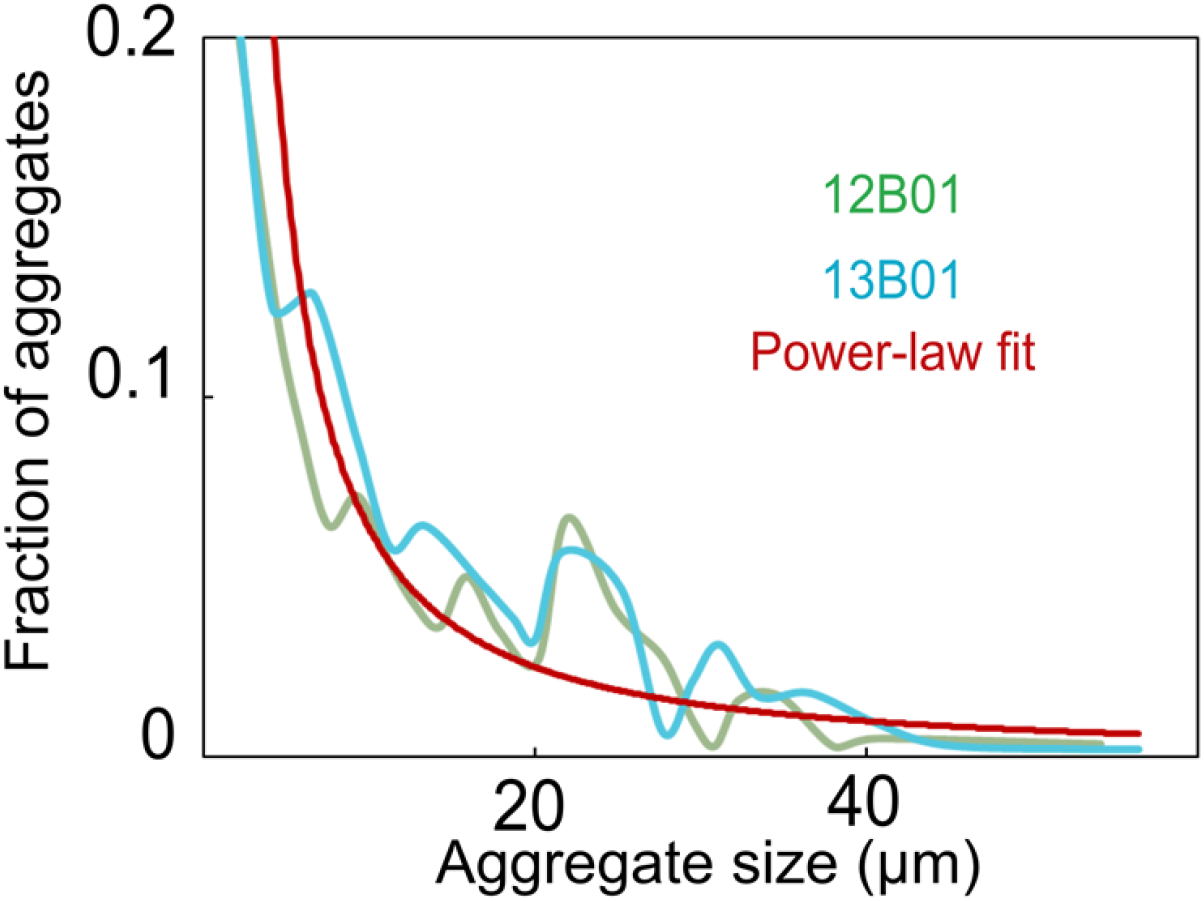
Quantification of aggregate size distributions for two Vibrio strains (12B01 and 13B01). Aggregates after 24 h from inoculation are measured. 250 aggregates are visualized uniform randomly over different fields of views of microscope.

